## Supplementary information for "Structural basis for human chondroitin sulfate chain polymerization"

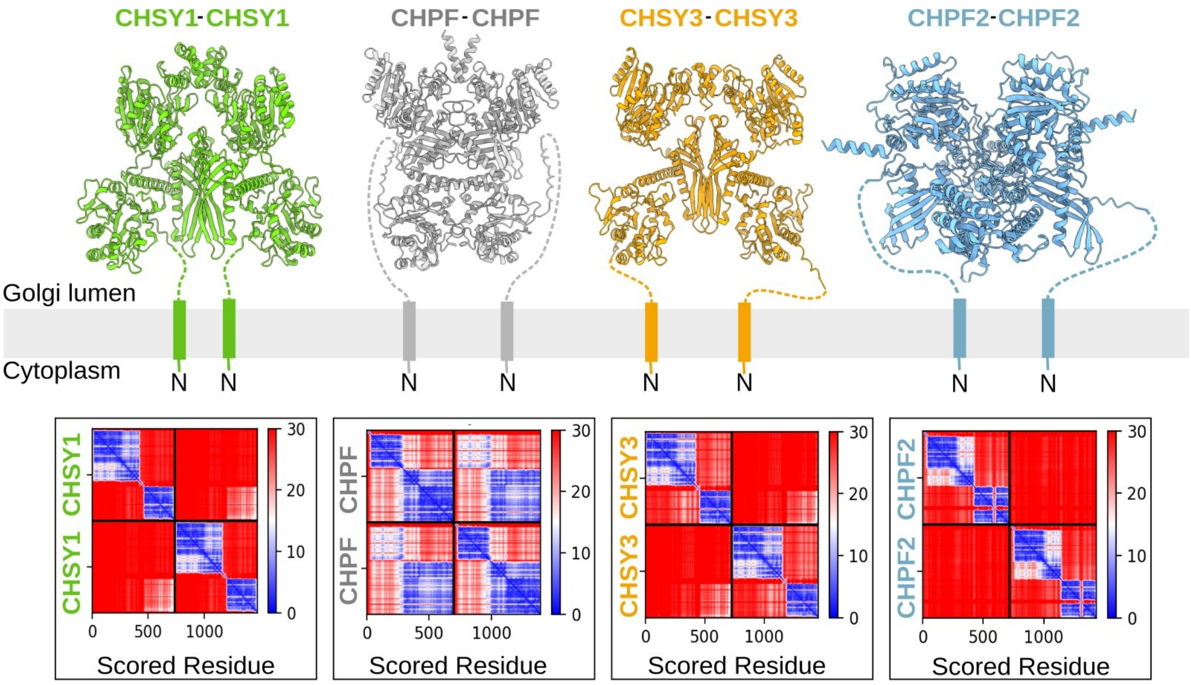


**Supplementary Fig. 1: CS synthase proteins do not seem to form homodimeric complexes.**
AlphaFold 2 predicted models for the four homodimeric CS polymerase complexes[1,2]. The models are shown in cartoon representation with CHSY1 in green, CHPF in grey, CHSY3 in orange and CHPF2 in light blue. The N-terminal anchoring helices and flexible stem regions (dotted lines), that were omitted during model prediction, were drawn by hand. The corresponding predicted align error matrices are shown below.

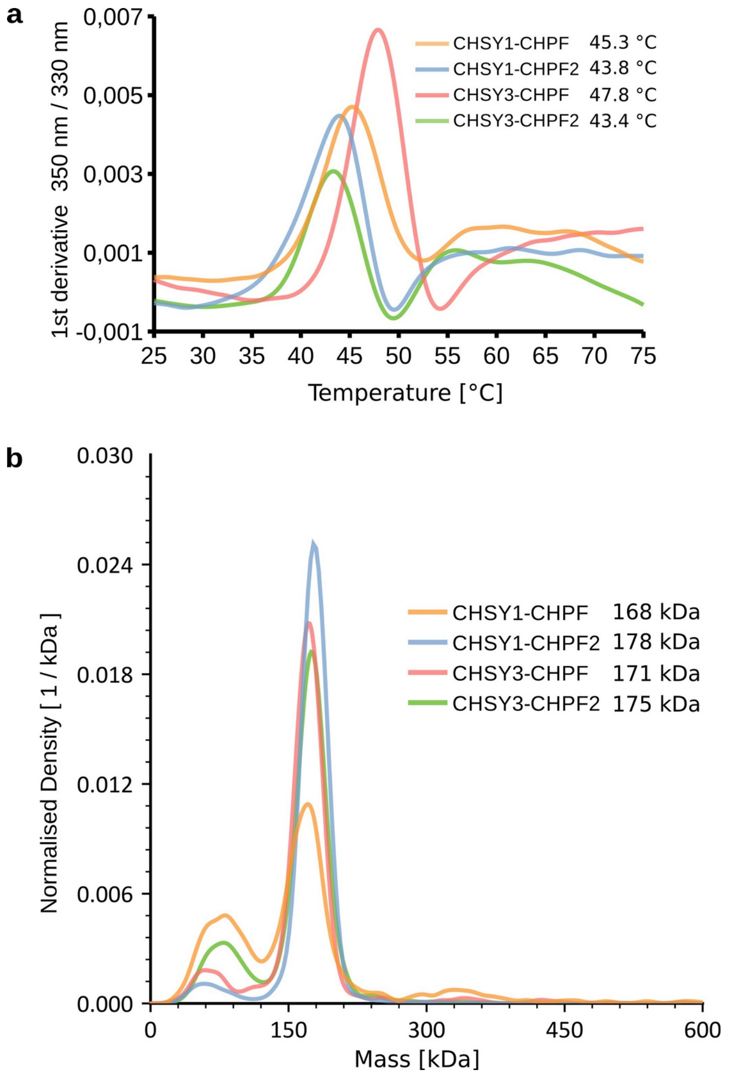


**Supplementary Fig. 2: Replicate measurements for wild-type CS polymerase complexes.**
**(a)** Duplicate measurement of melting temperature ($T_m$) of CS polymerase complexes using nano-differential scanning fluorimetry (nanoDSF). The obtained melting temperatures are indicated (see Fig. 2). **(b)** Duplicate measurement for mass photometry analysis of purified CS polymerase complexes with corresponding masses indicated alongside.

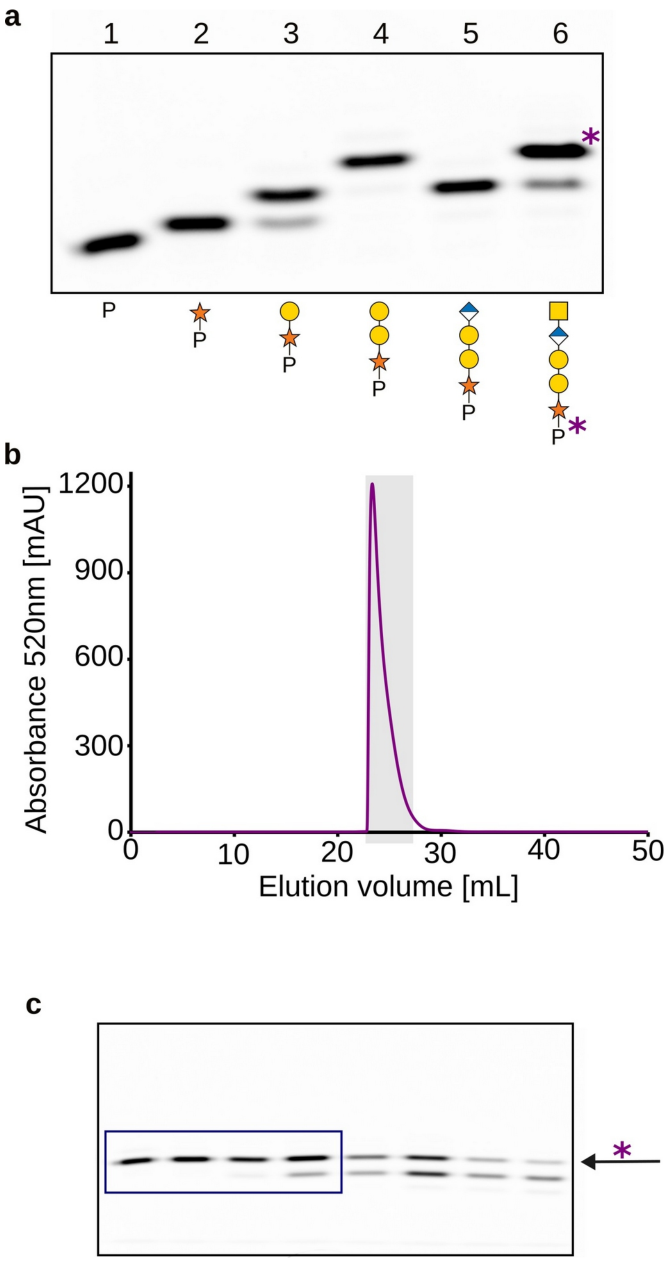


**Supplementary Fig. 3: Step-wise addition of pentsaccharide onto fluorescent CSF1 peptide.**
**(a)** FACE analysis of reaction products from chemo-enzymatic synthesis of pentasaccharide onto fluorescent CSF1 peptide. Lane 1 contains the fluorescent CSF1 peptide before glycan addition and lanes 2-6 show the peptide after mono-, di-, tri-, tetra and pentasaccharide addition, respectively. Each glycan addition is catalyzed by a distinct enzyme. Reaction products were visualized using a fluorescence imager and glycan addition can be followed by shifts in migration speed. Monosaccharide symbols follow the symbol nomenclature for glycans (SNFG) system[3]. **(b)** The pentasaccharide peptide generated in (a) was purified by size exclusion chromatography (SEC). (**c**) Peak fractions from SEC (highlighed in grey) were further analyzed by FACE. A purple asterisk indicates the pentasaccharide peptide product and fractions corresponding to lanes marked by a blue box were pooled.

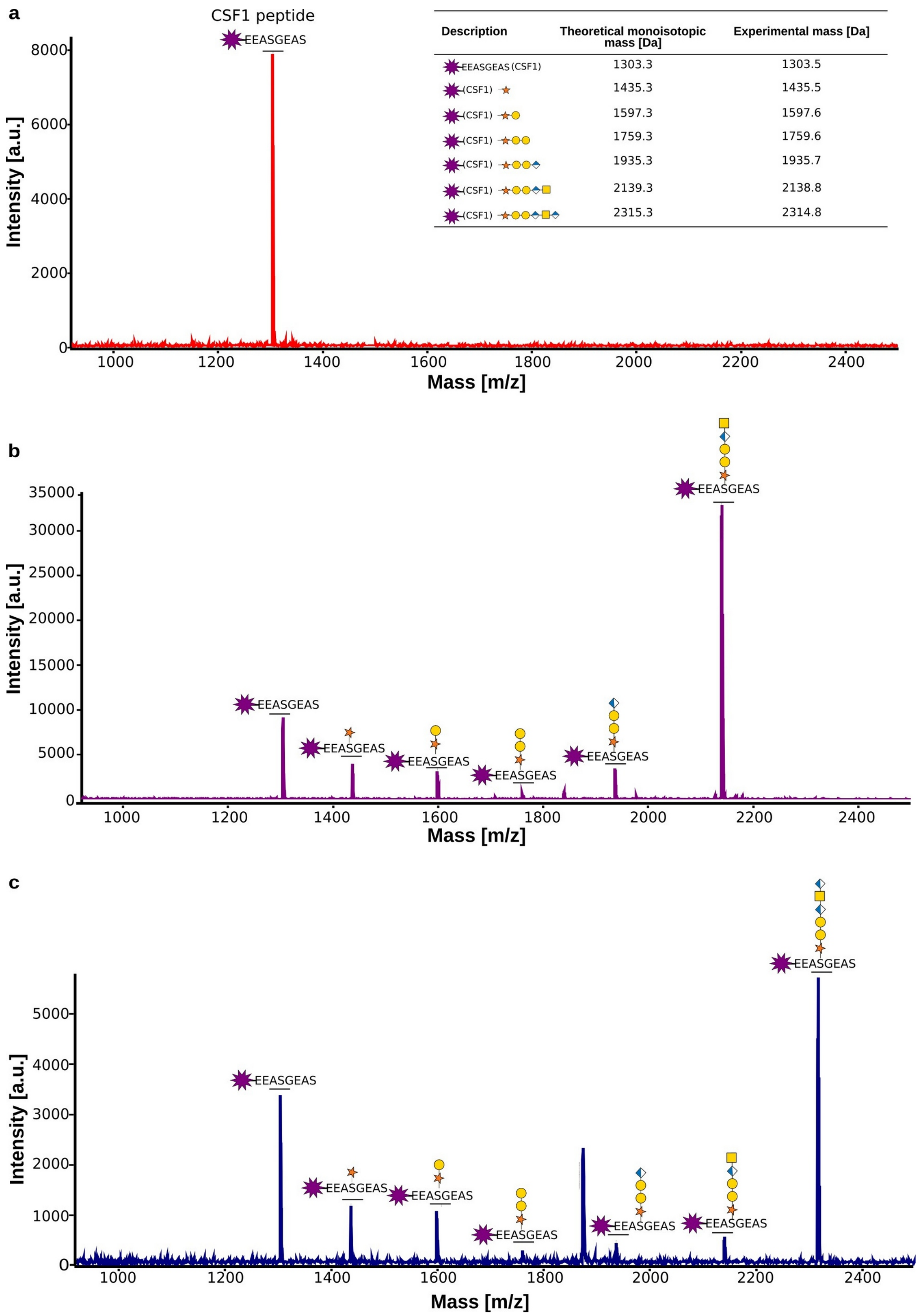


**Supplementary Fig. 4: Mass spectrometry analysis of Penta-CSF1 and Hexa-CSF1 peptides.**
**(a)** A synthetic fluorescent peptide (TAMRA-EEASGEAS) was derived from the chondroitin sulfate proteoglycan colony stimulating factor 1 (CSF1). This peptide was used as a control for MALDI-TOF analyses. **(b)** MS spectrum of purified Penta-CSF1 peptide carrying a pentasaccharide linker (Xyl-Gal-Gal-GlcA-GalNAc). Traces of reaction byproducts are present as well. **(c)** MS spectrum of purified Hexa-CSF1 peptide with a hexasaccharide linker (Xyl-Gal-Gal-GlcA-GalNAc-GlcA). The masses of the peptides are summarized in a table.

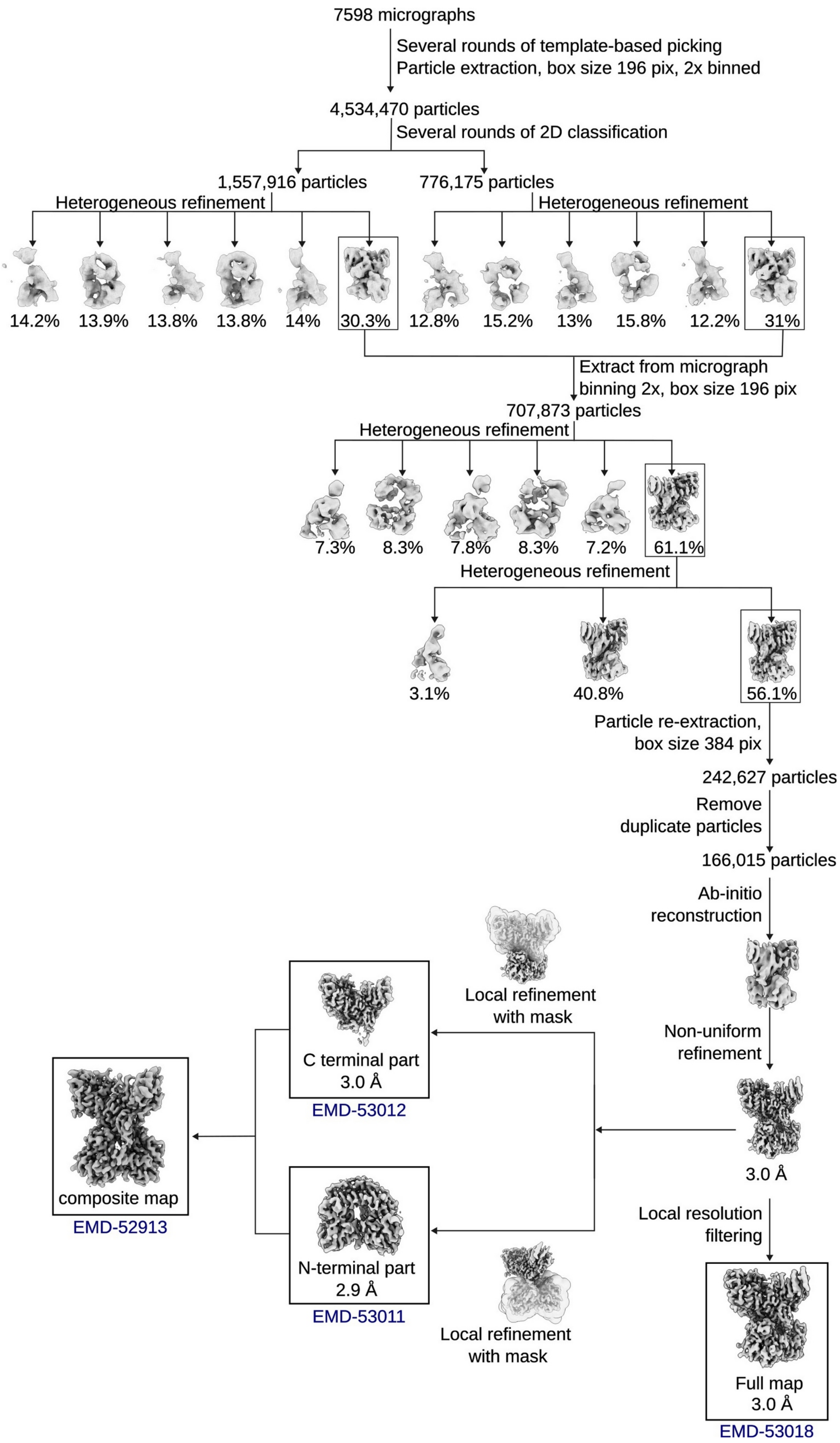


**Supplementary Fig. 5: Cryo-EM data processing flow chart.**
EM data was processed using cryoSPRAC v3.3.1 software. The full consensus map, the N- and C-terminal maps obtained from focused refinements and the generated composite map are accessible through the EMDB database. Corresponding accession codes are indicated.

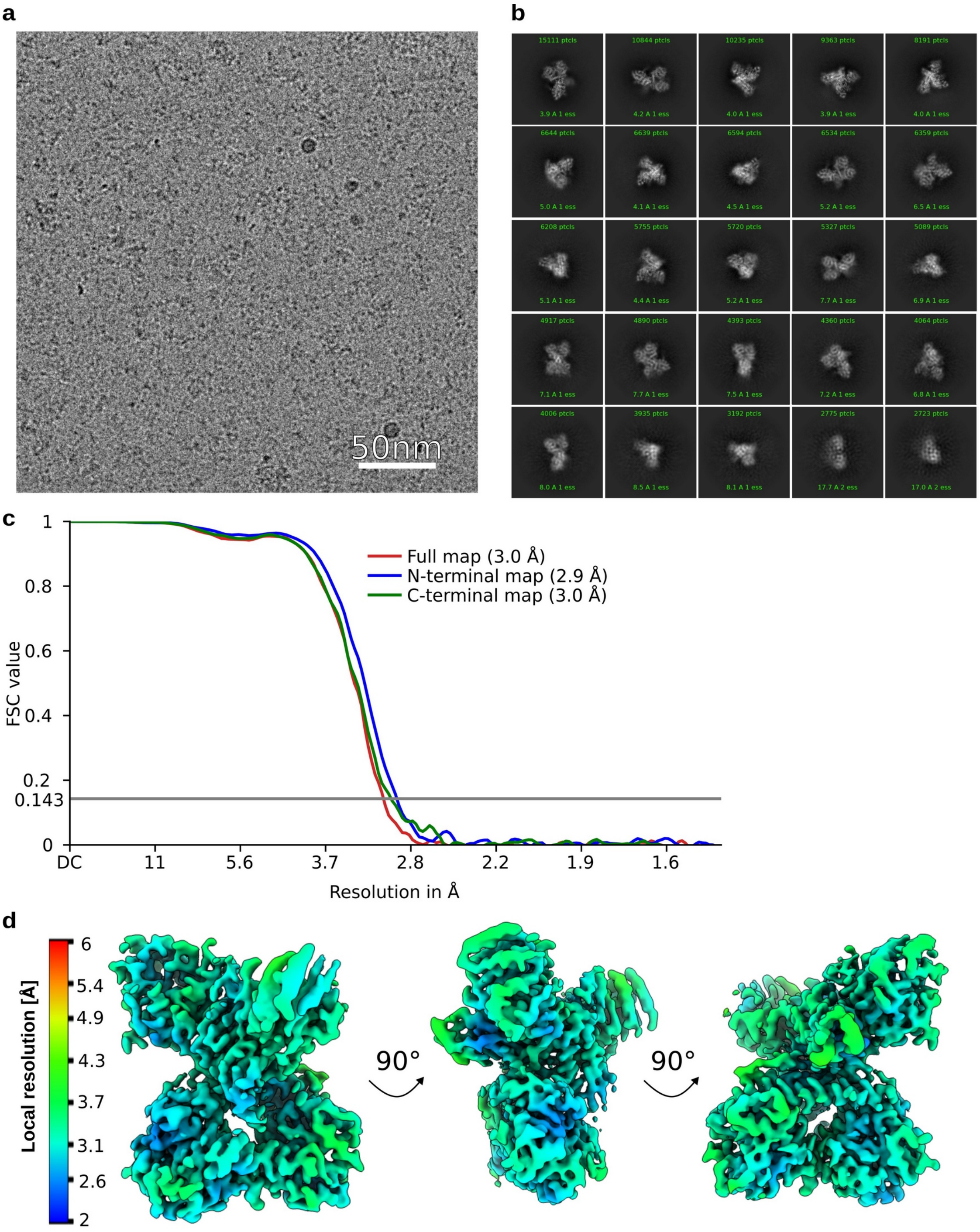


**Supplementary Fig. 6: EM data quality assessment.**

**(a)** Exemplary motion-corrected and dose-weighted micrograph. **(b)** 2D class averages of particles used for calculating final map. **(c)** Fourier shell correlation (FSC) curve indicating estimated resolutions based on the FSC = 0.143 criterion as generated by cryoSPARC v3.3.1. **(d)** Final local resolution-filtered EM map, colored by local resolution as estimated in phenix. The CS polymerase complex is shown from three orientations.

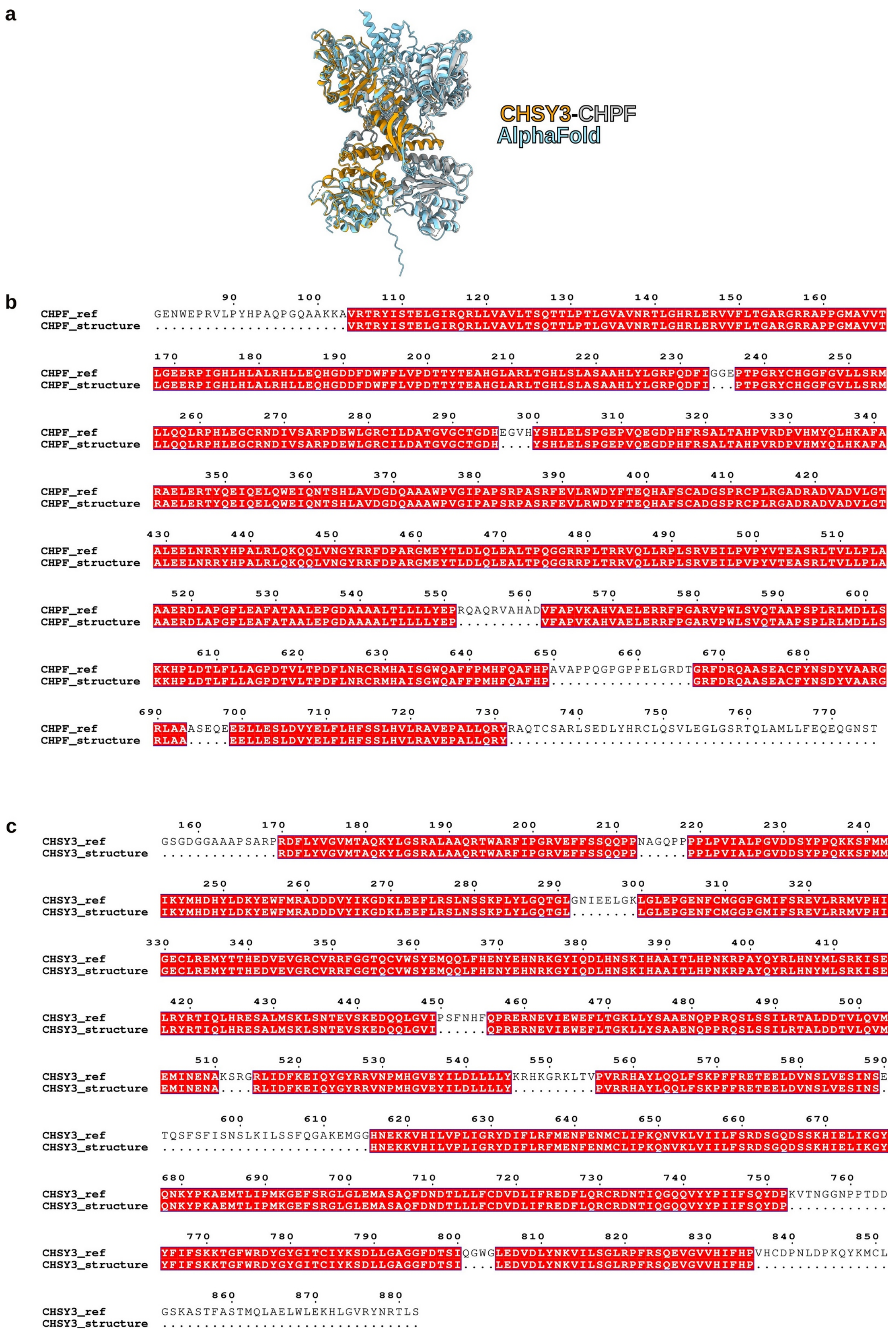


**Supplementary Fig. 7: Superposition of experimental and AlphaFold 2-predicted CHSY3-CHPF complex structure.**

**(a)** Cryo-EM structure of the CHSY3-CHPF complex, colored in orange and grey, and the model predicted using AlphaFold 2, colored in blue, were superimposed using the matchmaker command in ChimeraX[4]. The root mean square deviation (RMSD) between 522 pruned atom pairs (Cα) is 1.080 Å. **(b)** and **(c)** Alignment of the native protein sequences (ref) of CHPF and CHSY3 with the residues observed in the cryo-EM structure. The gaps reflect flexible loops and the N- and C-termini that were not visible.

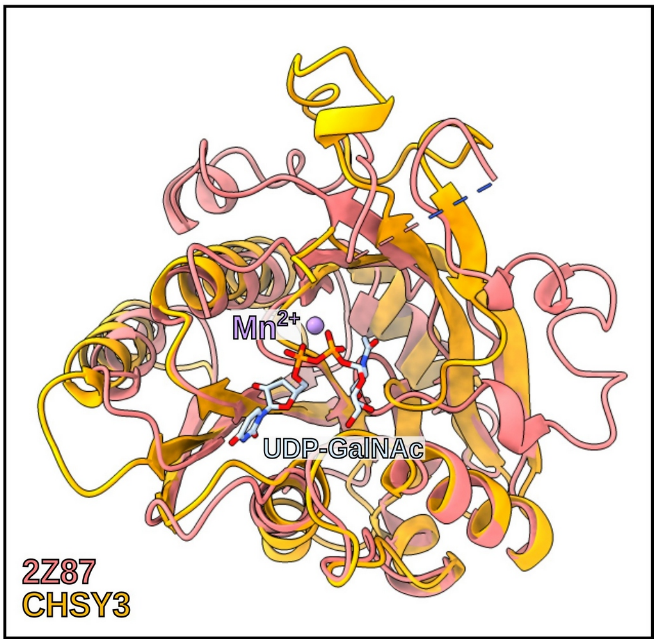


**Supplementary Fig. 8: Superposition of the GalNAc-T domain of CHSY3 with an *Escherichia coli* chondroitin polymerase.**
Superposition of the C-terminal GT domain of CHSY3 with the crystal structure of the close structural homolog Escherichia coli strain K4 chondroitin polymerase in complex with UDP-GalNAc (PDB ID: 2Z87). The two structures superimpose well with a root mean square deviation (RMSD) of 1.145 Å for 76 out of 286 residues (Cα).

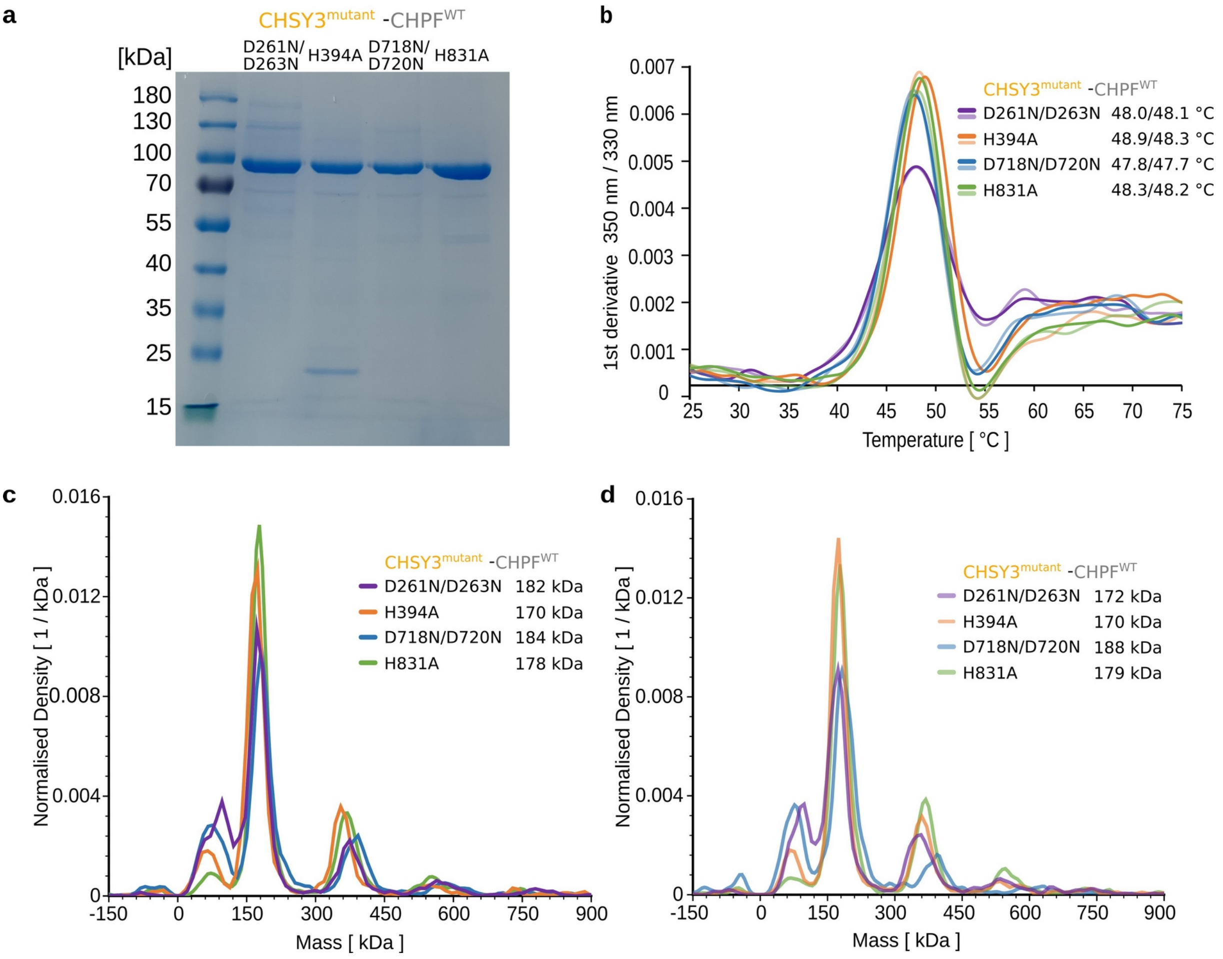


**Supplementary Fig. 9: Biophysical characterization of CHSY3-CHPF mutant complexes.**
**(a)** Coomassie-stained SDS-PAGE analysis of the four CHSY3 mutant-containing CHSY3-CHPF complexes. **(b)** Thermal stability of mutant CHSY3-CHPF complexes were determined by nanoDSF. Melting temperatures of duplicate measurements are indicated. **(c)** and **(d)** Mass photometry analysis of CHSY3 mutant-containing CHSY3-CHPF complexes performed as technical duplicates. The calculated molecular weights of complexes are indicated.

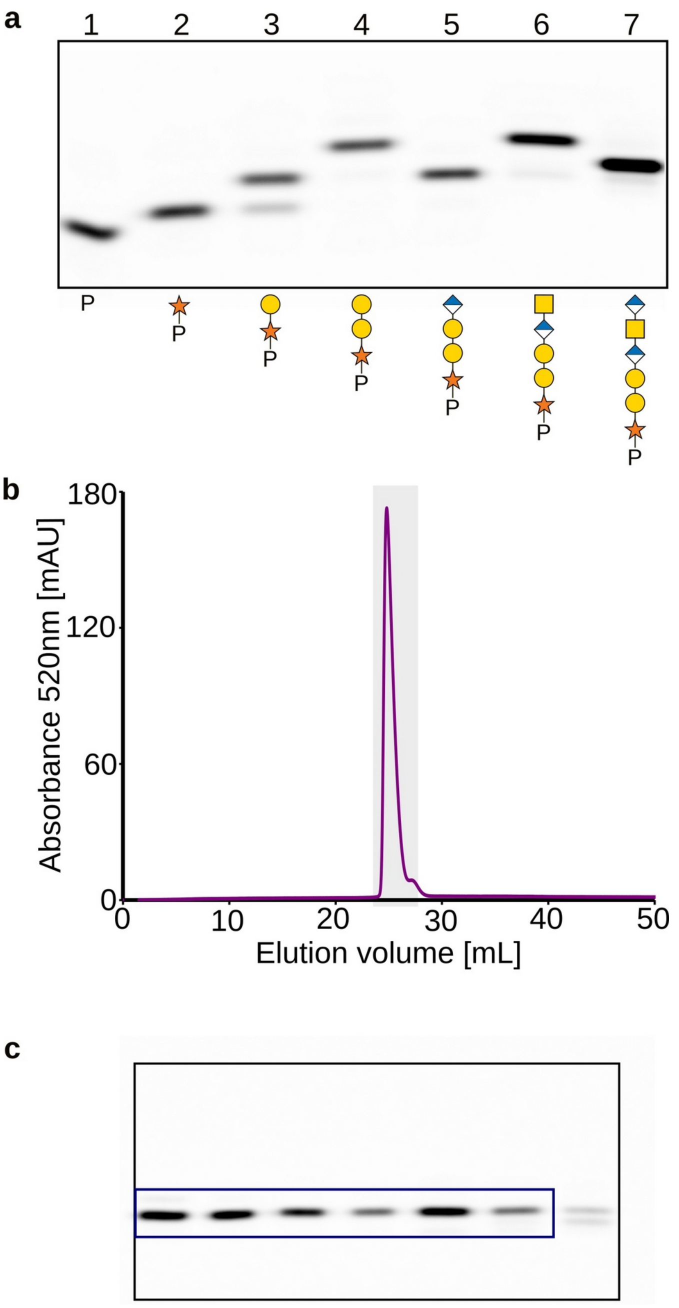


**Supplementary Fig. 10: Generation and purification of Hexa-CSF1 peptide.**

**(a)** Chemo-enzymatic synthesis of hexasaccharide onto CSF1 peptide was analyzed by FACE. Lane 1 contains the fluorescent CSF1 peptide before glycan addition and lanes 2-7 show the peptide after mono-, di-, tri-, tetra, penta- and hexasaccharide addition, respectively. Reaction products were visualized using a fluorescence imager and glycan addition can be followed by shifts in migration speed. Monosaccharide symbols follow the symbol nomenclature for glycans (SNFG) system[3]. **(b)** Hexa-CSF1 peptide was purified using SEC. (**c**) Peak fractions from SEC (highlighed in grey) were further analyzed by FACE. Hexasaccharide peptide containing fractions, marked by a blue box, were pooled.

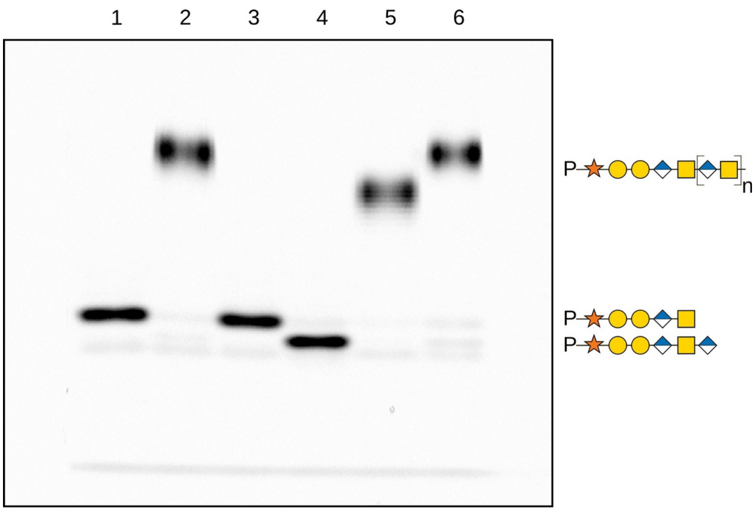


1: no enzyme

2: 125 nM WT

3: 125 nM D261N/D263N

4: 125 nM D718N/D720N

5: 62.5 nM D261N/D263N + 62.5 nM D718N/D720N

6: 125 nM D261N/D263N + 125 nM D718N/D720N

**Supplementary Fig. 11: Rescue experiment of mutant CHSY3-containing CS polymerase complexes.** Reactions containing wild-type (WT) or CHSY3 mutant-containing CHSY3/CHPF complexes, the Penta-CSF1 acceptor substrate, and both UDP-GalNAc and UDP-GlcA donor substrates were prepared. The ability of GlcA-T and GalNAc-T mutants to complement each other and thus restore chain elongation was tested. Reaction products were analyzed by FACE.

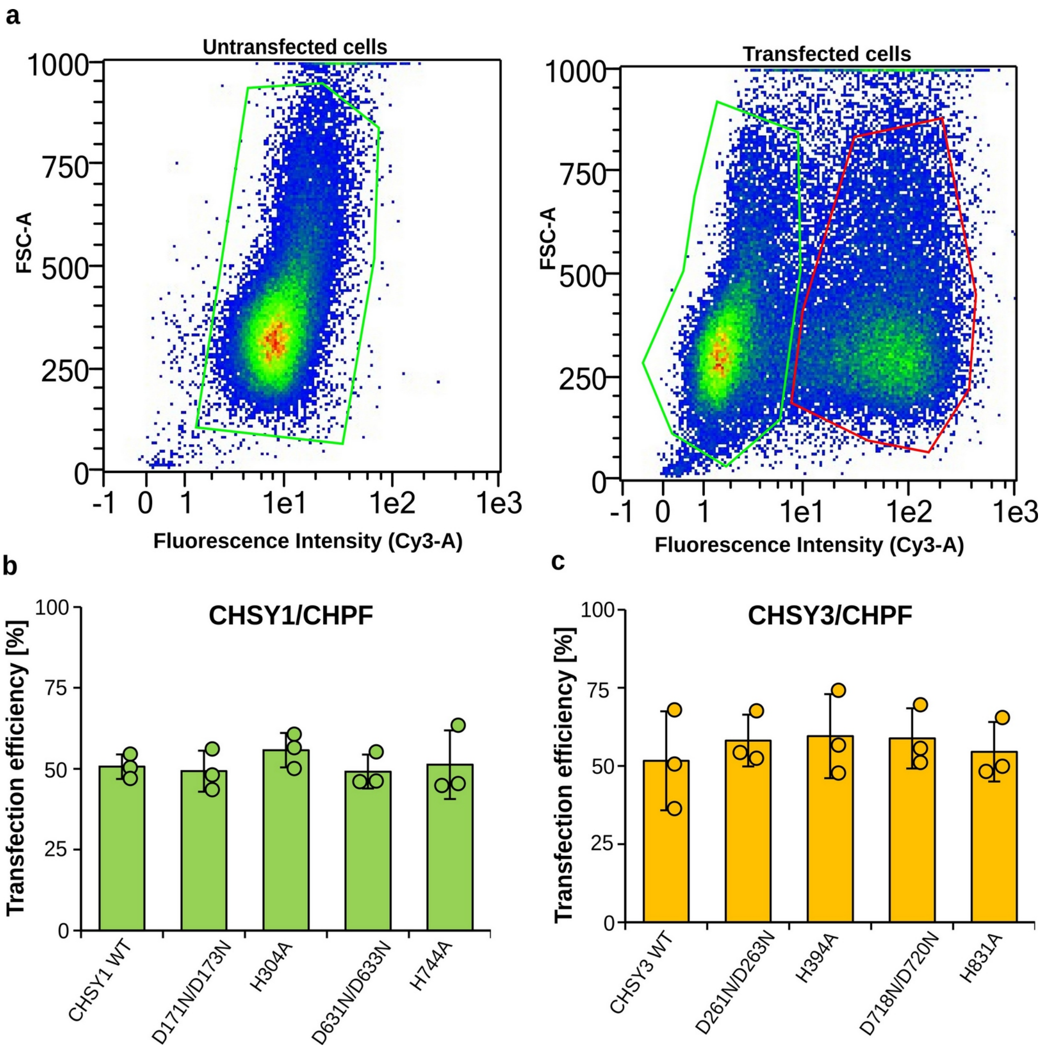


**Supplementary Fig. 12: Measurement of transfection efficiency by flow cytometry.**

**(a)** Exemplary scatter dot plots of flow cytometry experiments using an anti-FLAG primary antibody and a Cy3 secondary antibody. The left panel shows pattern of a cell population that was not transfected, and the right panel upon transfection with FLAG-CHSY3 and FLAG-CHPF encoding plasmids. Areas in the dot plot corresponding to untransfected and transfected cells are highlighted in green and red, respectively. **(b)** and **(c)** Bar plots showing the average transfection efficiency, calculated based on the number of successfully transfected cells (red box in a) in relation to total number of analyzed cells. Error bars show standard deviation from three independent experiments (n=3).

**a**

**Identity matrix [%]**

| | CHSY1 | CHSY3 | CHPF | CHPF2 |
|---|---|---|---|---|
| CHSY1 | 100 | | | |
| CHSY3 | 68.31 | 100 | | |
| CHPF | 23.73 | 24.59 | 100 | |
| CHPF2 | 24.61 | 24.2 | 59.15 | 100 |

**b**

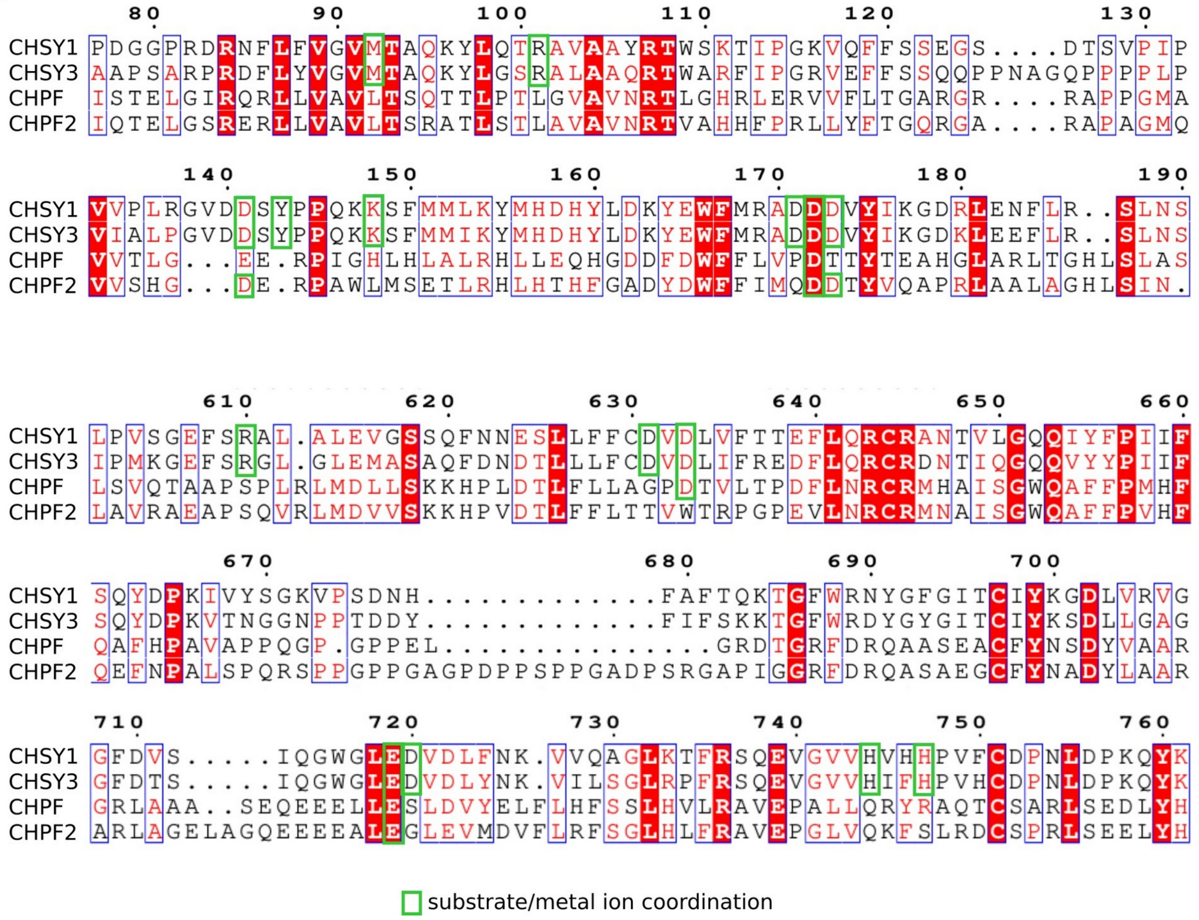


**Supplementary Fig. 13: Sequence analysis of human CS synthase proteins.**
**(a)** Sequence identity between full-length human CS synthase proteins CHSY1, CHSY3, CHPF and CHPF2 was calculated using Clustal Omega[5] and is displayed as an identity matrix. **(b)** Sequence alignment of catalytic regions of CHSY1, CHSY3, CHPF and CHPF2 was performed in ESPript 3.0[6]. Residues involved in UDP ligand and $Mn^{2+}$ coordination are highlighted in green.

**Supplementary Table 1: AlphaFold2 prediction scores**

Table summarizes the interface predicted template modeling (ipTM) values and predicted local distance difference test (pLDDT) values for AlphaFold2 predicted dimeric CS polymerase complexes.

| Complexes | pLDDT | ipTM |
|---|---|---|
| CHSY1-CHSY1 | 77 | 0.292 |
| CHPF-CHPF | 82.9 | 0.69 |
| CHSY3-CHSY3 | 77.8 | 0.3 |
| CHPF2-CHPF2 | 76.6 | 0.18 |
| CHSY1-CHPF | 89.9 | 0.921 |
| CHSY1-CHSY3 | 78.9 | 0.671 |
| CHSY1-CHPF2 | 89 | 0.911 |
| CHPF-CHSY3 | 89.9 | 0.919 |
| CHPF-CHPF2 | 81.8 | 0.364 |
| CHSY3-CHPF2 | 88.6 | 0.903 |

**Supplementary Table 2: EM data collection and refinement statistics**

| | |
|---|---|
| **Data collection** | |
| Microscope | Titan Krios CM02 G4 (Thermo Fisher Scientific) |
| Voltage | 300kV |
| Ditector | Falcon 4i |
| Energy filter | Selectris X energy filter cold FEG |
| Image format | .tiff |
| Magnification | 165,000 x |
| Defocus range (µm) | -2,1 to -0,5 |
| Total exposure dose ($e^-/Å^2$) | 50 |
| Dose rate ($e^-$/px/s) | 6.5 |
| Raw pixel size (Å) | 0.73 |
| Spherical aberration (mm) | 2.7 |
| Nr frames and fractions | 159 frames and 68 fractions |
| **Data processing** | |
| Software used for image processing | cryoSPARC v3.3.1 |
| Number of movies collected | 12,169 |
| Number of good micrographs | 7,598 |
| Final number of particles | 166,015 |
| Box size (pix) | 384 |
| Symmetry imposed | No |
| FSC threshold | 0.143 |
| Map resolution (Å) | 3.0 |
| N-ter map resolution (Å) | 2.9 |
| C-ter map resolution (Å) | 3.0 |
| Local filtering with B factor | -40 |
| Map visualizing software | ChimeraX |
| **Model building and refinement** | |
| PDB accession ID | 9Q8Z |
| Initial model used | Alphafold 2 model |
| Model composition<br>Non-hydrogen atoms<br>Protein residues<br>Ligands | <br>9612<br>1179<br>2 MN; 1 UDP; 3 NAG |
| Root mean square deviations<br>Bond lengths (Å)<br>Bond angles (°) | <br>0.003<br>0.545 |
| Ramachandran plot<br>Favored (%)<br>Allowed (%)<br>Disallowed (%) | <br>95.82<br>4.18<br>0 |
| Validation<br>Molprobity score<br>Clashscore<br>Poor rotamers (%) | <br>1.8<br>9.79<br>0 |

**Supplementary Table 3: Primer for site-directed mutagenesis PCR**

| Mutation | Forward primer (5’- 3’ sequence) | Reverse primer (5’- 3’ sequence) |
|---|---|---|
| **CHSY1 D171N/D173N** | GT GCC AAC GAC AAT GTC TAT ATT AAG GGC GAT AGA CTT GAA | GAC ATT GTC GTT GGC AC GCA TGA ACC ACT CGT ATT TAT CC |
| **CHSY1 H304A** | T ACT CTT GCT CCA AAT A AGA ATC CCC CAT ATC AAT ATC GA | T ATT TGG AGC AAG AGT A ATC GCC TGA TGT ATC TTG GAA |
| **CHSY1 D631N/D633N** | TTC TTT TGT AAT GTA AAT TTG GTC TTC ACA ACT GAG TTT CTG C | CAA ATT TAC ATT ACA AAA GAA CAA GAG GCT CTC GTT ATT GAA |
| **CHSY1 H744A** | C GTG GTT GCT GTT CAT CAC CCA GTG TTC TGC GAC C | ATG AAC AGC AAC CAC G CCC ACC TCT TGA CTT CTG AAT GTT |
| **CHSY3 D261N/D263N** | GG GCA AAT GAT AAC GTGTAT ATA AAG GGC GAC AAG CTT GA | CAC GTT ATC ATT TGC CC TCA TAA ACC ATT CGT ACT TAT CGA G |
| **CHSY3 H394A** | ACG CTG GCC CCT AAT AAG CGC CCA GCC TAT CAG TA | ATT AGG GGC CAG CGT AAT TGC GGC GTG AAT CTT AG |
| **CHSY3 D718N/D720N** | TTC TGC AAC GTA AAT CTG ATC TTT CGG GAA GAC TTC TTG C | CAG ATT TAC GTT GCA GAA CAG AAG CAG GGT ATC GTT ATC G |
| **CHSY3 H831A** | GT GTA GTT GCC ATC TTT C ACC CCG TGC ACT GCG AC | GAA AGA TGG CAA CTA CAC CCA CCT CTT GGG ACC GA |
